# Postpartum fluoxetine increases maternal hippocampal IL-1β and decreased plasma tryptophan: clues for efficacy

**DOI:** 10.1101/2020.02.22.960021

**Authors:** Wansu Qiu, Paula Duarte-Guterman, Rand S. Eid, Kimberly A. Go, Yvonne Lamers, Liisa A. M. Galea

## Abstract

Perinatal depression (PND) affects approximately 15% of women, and *de novo* postpartum depression affects approximately 40% of PND cases. Selective serotonin reuptake inhibitors (SSRIs) are a common class of antidepressants prescribed to treat PND. However, the safety and efficacy of SSRIs have been questioned in both clinical and preclinical research. Here, using a preclinical rodent model of postpartum depression, we aim to better understand neuroinflammatory cytokines and tryptophan mechanisms that may be related to SSRIs efficacy. Rodent dams were treated with high corticosterone (CORT; 40mg/kg, s.c.) for 21 days in the postpartum period to simulate depressive-like behaviors in the late postpartum period. Concurrently, a subset of dams was treated with the SSRI, fluoxetine (FLX; 10mg/kg, s.c.), in the postpartum period. We showed, consistent with previous studies, that although maternal FLX treatment prevented CORT-induced disturbances in maternal care behavior during the early postpartum, it failed to prevent the expression of CORT-induced depressive-like behavior in the late postpartum. Furthermore, FLX treatment, regardless of CORT treatment, increased maternal hippocampal IL-1β and decreased maternal plasma tryptophan levels, plasma tryptophan, 4’-pyridoxic acid, and pyridoxal concentrations. Maternal CORT treatment reduced maternal hippocampal TNF-α and IFN-γ levels. Our work suggests that the limited efficacy of FLX in the late postpartum may be associated with elevated levels of the proinflammatory cytokine IL-1β in the maternal hippocampus, decreased plasma tryptophan concentration, and changes in vitamin B6 dependent tryptophan-kynurenine pathway. These findings suggest novel pathways for improving SSRI efficacy in alleviating perinatal depression.

**HIGHLIGHTS:**

- Postpartum fluoxetine (FLX) increased interleukin-1β levels in hippocampus
- Postpartum corticosterone (CORT) decreased TNF-α and IFN-γ in the hippocampus
- Postpartum FLX did not prevent CORT-induced depressive-like behavior
- Postpartum FLX prevented CORT-induced changes in maternal behavior
- Postpartum FLX decreased plasma tryptophan, 4’-pyridoxic acid, and pyridoxal levels

## 1. INTRODUCTION

Perinatal depression (PND) affects approximately 15% of women within the general population (O’Hara and Swain, 1996; Shorey et al., 2018), and depression onset can occur *de novo* during pregnancy or in the postpartum. Untreated PND can have severe and long-lasting effects on the mother and her children (Beck, 1999; Grace et al., 2003; Weissman et al., 2006). However, the efficacy and safety of maternal antidepressant use during pregnancy and/or postpartum has been challenged in both humans and animal models (Oberlander et al., 2009; Sharp et al., 2010; De Crescenzo et al., 2014; Suri et al., 2017; Workman et al., 2016; Gobinath et al., 2018). Thus, it is important to determine factors during the perinatal period that may contribute to reduced drug efficacy during the perinatal period.

One of the strongest predictors of PND is stressful life events (Robertson et al., 2004). Additionally, in response to stress, a subset of women with PND show dysregulation of the hypothalamic-pituitary-adrenal (HPA) axis (Jolley et al., 2007; Glynn et al., 2013; Osborne et al., 2018) and higher evening salivary cortisol, the main stress hormone, when compared to healthy controls (Iliadis et al., 2015; Osborne et al., 2018). Animal models of depression often utilize stress as an inducer of depressive-like phenotypes (Krishnan and Nestler, 2008) and models of PND use either gestational stress (Hillerer et al., 2011; Pawluski et al., 2011; Haim et al., 2014) or postpartum corticosterone (CORT) (Brummelte et al., 2006; Brummelte and Galea, 2010; Workman et al., 2013). Both models of PND recapitulate different etiologies and ontogeny timelines of depressive-like behaviors, which mirrors the heterogeneity of PND (Postpartum Depression: Action Towards Causes and Treatment (PACT) Consortium, 2015; Galea and Frokjaer, 2019). Gestational stress models depression that would occur *de novo* during pregnancy, whereas postpartum corticosterone models depression that occurs *de novo* in the postpartum only. Although depression during pregnancy is one of the greatest predictors of postpartum depression, at least 40% of women with PND will develop depression for the first time during the postpartum period (Munk-Olsen et al., 2006; Wisner et al., 2013).

Intriguingly, in rodent models, the effects of gestational stress on depressive-like endophenotypes can be restored with pharmacological antidepressants, such as selective serotonin reuptake inhibitors (SSRIs) (Gemmel et al., 2016; Haim et al., 2016), while, there is less efficacy of SSRIs during the postpartum using the CORT-induced model (Workman et al., 2016; Gobinath et al., 2018). In the CORT-induced model of postpartum depression, SSRI treatment is effective at rescuing disrupted maternal care in the short term (days 2-8 after parturition) but is ineffective in alleviating passive-coping or reduced hippocampal neurogenesis in the long-term (days 23-24 after parturition) (Workman et al., 2016; Gobinath et al., 2018). Based on these data, it is important to better understand the possible mechanisms that may be related to the lack of efficacy of SSRIs during the late postpartum period, and we chose to examine two potential factors: neuroinflammation and plasma metabolites of the vitamin B6 dependent tryptophan-kynurenine pathway.

Inflammation is associated with major depressive disorder (MDD) as patients diagnosed with MDD show higher proinflammatory cytokines, including tumour necrosis factor (TNF)-α, interleukin (IL)-6, and IL-1β (Dowlati et al., 2010; Haapakoski et al., 2015; Howren et al., 2009). In addition, pregnancy and the postpartum also induce changes in the immune system in the hippocampus (Haim et al., 2017; Sherer et al., 2017) and in blood (Eid et al., 2019; reviewed in Duarte-Guterman et al., 2019). Elevated levels of peripheral proinflammatory immune markers, particularly the cytokines IL-1β, interferon-gamma (IFN-γ) and IL-6, are reported in women endorsing PND (Groer and Morgan, 2007; Corwin et al., 2008) or associated with depression scores in the perinatal period (Maes et al., 2000), implicating the immune system in PND. These associations and natural changes in the immune system during the perinatal period may offer clues as to the lack of SSRI efficacy during this period. Indeed, treatment non-responsive patients either show no reduction in cytokine levels or continued elevation of cytokine levels (Syed et al., 2018), while depressed patients in remission show reductions in peripheral cytokine levels in response to SSRI treatment (Hannestad et al., 2011; Syed et al., 2018), Moreover, Haroon and colleagues (2018) found an association between higher levels of cytokines TNF and IL-6, and the number of failed treatment trials (Haroon et al., 2018). These findings suggest that inflammation signalling may give us clues as to the lack of SSRI efficacy in the postpartum period.

Another possible mechanism associated with antidepressant efficacy is tryptophan metabolism and the vitamin B6 dependent tryptophan-kynurenine pathway. Elevated tryptophan metabolism as indicated by lower tryptophan concentrations (Hughes et al., 2012; Myint et al., 2013; Ogawa et al., 2014; Doolin et al., 2018, Pu et al., 2020) and higher metabolite concentrations are associated with MDD (Myint et al., 2007; Savitz 2016; Ogyu et al., 2018; Pu et al., 2020). Furthermore, inflammation can increase tryptophan metabolism (Myint et al., 2013), and thus influence serotonin synthesis, and this relationship is one proposed mechanism through which inflammation can lead to depression onset (Myint et al., 2013). Importantly, MDD individuals who are non-responders to SSRI treatment had higher metabolite markers of tryptophan metabolism at baseline (Sun et al., 2020). Interestingly, the tryptophan-kynurenine pathway regulates hippocampal neurogenesis via IL-1β (Zunszain et al., 2012), and neurogenesis is decreased in CORT-induced model of postpartum depression (Brummelte and Galea, 2010; Workman et al., 2016; Gobinath et al., 2018). Thus, levels of tryptophan metabolism may be a modulator of SSRI efficacy.

Two brain regions, the hippocampus and the frontal cortex, are both highly associated with depression (Fanselow and Dong, 2010; Koenigs and Grafman, 2009). For example, a meta-analysis indicates that people diagnosed with MDD for at least two years present with smaller hippocampal volume (McKinnon et al., 2009). In a separate meta-analysis of resting-state functional connectivity studies, MDD was found to be characterized by hypoconnectivity in frontoparietal regions important for regulation of attention and emotion (Kaiser et al., 2015). Interestingly, the postpartum period is associated with reduced gray matter in both the hippocampus and various regions in the prefrontal cortex (Hoekzema et al., 2017). Models of PND result in decreased spine density in dams in the prelimbic medial frontal cortex (Leuner et al., 2014; Haim et al., 2016) and/or reduced neurogenesis in the hippocampus (Pawluski and Galea, 2007; Workman et al., 2016; Gobinath et al., 2018). Thus, it is clear that the pathoetiology of depression involves disruptions to both the frontal cortex and hippocampus, two areas that are examined in this paper.

Here, we sought to characterize markers of inflammation in the maternal hippocampus and frontal cortex, and plasma tryptophan metabolism markers in a CORT-induced model of postpartum depression and SSRI treatment. We hypothesized that CORT treatment would induce depressive-like endophenotypes, including disruptions in maternal care behavior and increased passive-coping behavior. Furthermore, we expected that the SSRI, fluoxetine (FLX), will show limited efficacy at reducing CORT-induce depressive-like endophenotypes. We also hypothesized that FLX treatment would alter markers of neuroinflammation and tryptophan metabolism, which could be related to the limited efficacy.

## 2. METHODS

### 2.1 Animals

Thirty-seven adult female Sprague-Dawley rats and six male rats (2.5 months old) were purchased from Charles River (Montreal, QC, Canada). All animals were initially housed in same-sex pairs upon arrival in polycarbonate cages (24 × 16 × 46 cm) with aspen chip bedding. Males and females were housed in separate colony rooms. All animals were maintained on a 12 h: 12 h light/dark cycle (lights on at 07:00) and given standard rat chow (Jamieson’s Pet Food Distributors Ltd, Delta, BC, Canada) and autoclaved tap water *ad libitum*. All animals were allowed to acclimatize to the facilities for seven days and handled for five days prior to the onset of the experiment. Females were randomly assigned to one of four experimental groups (described below), and males were utilized only as breeders. All protocols were in accordance with ethical guidelines set by the Canadian Council for Animal Care and were approved by the University of British Columbia Animal Care Committee. For an overview of experimental procedures, refer to Figure 1.

**Figure 1.**
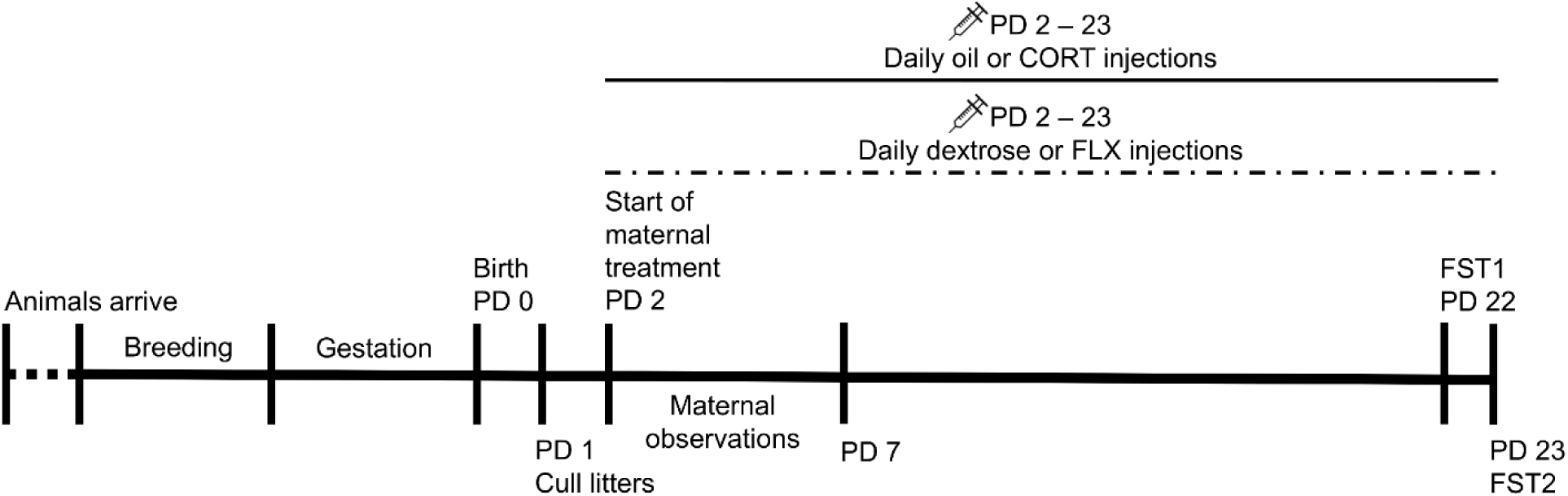
Timeline of experiment. Briefly, animals were bred within the animal facility and kept undisturbed throughout gestation. Date of birth was designated postnatal day (PD) 0. Upon PD2, animals received daily injections of either corticosterone (CORT) or oil vehicle and fluoxetine (FLX) or dextrose vehicle for 21 days. Maternal care observations occurred during PD2-7. Forced-swim test (FST) was conducted on PD22 and PD23. Pups remained with the dams until experimental endpoint.

### 2.2 Breeding and culling

Breeding and culling protocols were followed as previously described (Workman et al., 2013). Briefly, males were single-housed for 24 hrs prior to breeding. Upon breeding, two females were co-housed with a single male overnight. Vaginal lavage samples were collected the following morning and females were deemed pregnant if sperm was detected (considered gestational date (GD)1). Females were single-housed on GD1 and left undisturbed for the duration of gestation in a separate colony room, except for weekly cage-changing and weighing.

All litters were culled randomly on postpartum day 1 (PD1, the following day after birth of the pups) to 5 females and 5 males per litter. A total of two dams received cross-fostered pups from aged-matched groups because original litters did not meet the requirement of 5 females and 5 males.

### 2.3 Drug preparations and treatments

All treatments were administered as previously described (Gobinath et al., 2018). All animals received daily injections beginning from PD2 to PD23 between 08:00-10:00. Dams received corticosterone (CORT; 40mg/kg, s.c.; Sigma-Aldrich, St. Louis, MO, USA) or its vehicle (OIL; sesame oil + 10% EtOH, 1ml/kg, s.c.) and either fluoxetine (FLX; 10mg/kg, s.c.; Sequoia Research Products, Pangbourne, UK) or its vehicle, dextrose (DXT; 5% dextrose in sterile water, 1ml/kg, s.c.). Thus, all dams were randomly assigned to one out of four treatment groups (OIL+DXT, OIL+FLX, CORT+DXT, and CORT+FLX). FLX and an emulsion of CORT were prepared every three days. This dose of CORT has reliably produced a depressive-like phenotype in dams with reduced maternal care, reduced neurogenesis in the hippocampus, and increased immobility in the forced swim test (Brummelte et al., 2006; Gobinath et al., 2018).

### 2.4 Maternal care behavior observations

Maternal care behavior observations were conducted twice per day (morning and afternoon) for six days during PD2-7. Observations occurred at least 2hrs after injections or after the previous observation. Each observation lasted for 10 minutes, and the following maternal care behaviors were scored: (a) nursing, which included arched-back nursing (arched-back, actively nursing pups), blanket nursing (in nursing position but resting on-top of pups), and passive nursing (lying on the side and nursing pups); (b) licking (licking pups while not nursing any other pups), (c) self-grooming (not engaging with pups, and self-grooming), (d) nest building (actively building nest), and (e) time spent off-nest (off-nest and not engaging with pups).

### 2.5 Forced-swim test

Dams were tested in the forced-swim test (FST) on PD22 (FST1) for 15min and PD23 (FST2) for 5min to assess the effects of CORT and/or FLX on passive-coping behavior as previously described (Workman et al., 2016; Gobinath et al., 2018). FST is a reliable measure of antidepressant efficacy (Lucki, 1997; Cryan et al., 2002). All behavioral testing occurred at least 2 hours after injections in a separate behavioral room. Briefly, all dams were placed into a water tank containing warm, clean water (24-26°C) in dim light. Behaviors were videotaped and scored by a blinded scorer using BEST Collection Software (Educational Consulting, Inc, Hobe Sound, FL, USA). During FST1, pups were left undisturbed outside of behavioral room and weaned during FST2. Three specific behaviors were coded for: (1) swimming (animals were actively moving their limbs), (2) immobility (immobile with only movements necessary to stay floating), and (3) climbing along tank walls (animals were actively moving limbs along the tank walls, attempting to climb).

### 2.6 Estrous cycle cytology

Vaginal lavage samples were taken prior to FST2 and were counterstained using cresyl-violet solution to be assessed later for estrous cycle stages based on McLean et al. (2012) and Cora et al. (2015).

### 2.7 Blood and tissue collection

Immediately after FST2, all animals were euthanized by rapid decapitation, and trunk blood was collected within three minutes of touching the cage. Plasma was collected into cold EDTA coated tubes and centrifuges 4h later for 10min at 4°C. Brains and adrenal glands were extracted on a cold surface. Hippocampus (0.60mm above bregma to −7.20mm below bregma) and frontal cortex (6.12mm to 0.60mm above bregma) regions were microdissected over ice and flash-frozen immediately after extraction on dry ice and stored at −80°C until assayed for protein and cytokine levels. Adrenal glands were collected and weighed.

### 2.8 Tissue homogenization

Following previous protocols (Bodnar et al., 2017; Eid et al., 2019), briefly, all tissues were homogenized in cold lysis buffer using the Omni Bead Ruptor 24 (Omni International. Kennesaw, GA, USA). Immediately after homogenization, all tissue samples were centrifuged at 4°C at 13.2 rpm for 20min. Supernatant was then collected and stored in −80°C for protein and cytokine analysis.

### 2.9 Cytokine and protein quantification

Cytokine concentrations were measured as previously described (Galea et al., 2018; Eid et al., 2019) using a multiplex electrochemiluminescence immunoassay kit (V-PLEX Proinflammatory Panel 2, Rat) from Meso Scale Discovery (Rockville, MD, USA) used according to manufacturer’s instructions. Samples were run in duplicates to quantify in multiplex in each sample: interferon-gamma (IFN-γ), interleukin-1-beta (IL-1β), interleukin-4 (IL-4), interleukin-5 (IL-5), interleukin-6 (IL-6), tumour necrosis factor-alpha (TNF-α) and chemokine (C-X-C motif) ligand 1 (CXCL1). Plates were read using a Sector Imager 2400 (Meso Scale Discovery), and data analyses were conducted using the Discovery Workbench 4.0 software (Meso Scale Discovery). Lower limits of detection (LLOD) were as follows (pg/mL) for HPC: INF-γ: 0.568, IL-1β: 2.31, IL-4: 0.399, IL-5: 9.35, IL-6, 5.78, IL-10: 2.53, IL-13: 1.08, TNF-α: 1.36, and CXCL1: 0.357; and for PFC: INF-γ: 0.179, IL-1β: 2.79, IL-4: 0.143, IL-5: 4.08, IL-6, 8.67, IL-10: 0.603, IL-13: 0.429, TNF-α: 0.331, and CXCL1: 0.467. Seven sample duplicates with high intra-assay coefficients of variations (>15%) were eliminated from the final analyses. No values were below LLOD.

Total protein levels were quantified in tissue homogenates using the Pierce Microplate BCA Protein Assay Kit in triplicates following the manufacturer’s protocol (Thermo Scientific Pierce Protein Biology, Thermo Fisher Scientific, Waltham, MA, USA). All cytokine levels were normalized by total protein levels and reported as pg cytokine/mg of protein.

### 2.10 Plasma pyridoxic acid, pyridoxal, and tryptophan analyses

We quantified vitamin B6 and related metabolites because vitamin B6 in its coenzyme form pyridoxal 5’-phosphate is a critical coenzyme in tryptophan metabolism (Bender et al., 1990). Plasma 4’-pyridoxic acid, pyridoxal, pyridoxal 5’-phosphate, and tryptophan concentrations were quantified using isotope dilution liquid chromatography coupled with tandem mass spectrometry-based on the method developed modified by Midttun et al. (2005; 2009) with modifications. The intra- and interassay CVs for these assays were all >7%

### 2.11 Statistical analyses

All data (maternal behaviors, relative adrenal mass, FST behaviors, tryptophan metabolites, and cytokines) were analyzed using two-way analysis of variance (ANOVA) with CORT (CORT or OIL) and FLX (FLX or DXT) as between-subject factors unless otherwise specified. Relative adrenal mass was calculated as adrenal mass/body mass. Effect sizes are reported for significant effects as η_p_^2^ or Cohen’s d. Post hoc comparisons used Newman-Keuls. *A priori* comparisons were subjected to Bonferroni correction. As an exploratory approach, principal component analyses (PCA) were conducted to determine networks (components) that best explain cytokines level variances within a specific brain region (HPC or PFC). Principal component scores for individual sample data for the first two components generated were then analyzed using ANOVA models to derive information regarding the amount of variance accounted for by cytokine profiles within the hippocampal or frontal cortex data set (Eid et al., 2019). All ANOVA and correlation data analyses were analyzed using Statistica (v. 9, StatSoft, Inc., Tulsa, OK, USA). PCA data analyses were conducted using R (3.4.3) statistical analysis software with the FactoMineR package. Effects were considered statistically significant if *p* ≤ 0.05 and trends are discussed if *p* ≤ 0.10. A priori comparison were based on previous literature showing that postpartum CORT reduces nursing and increases time spent off the nest, while postpartum FLX restores these deficits (Workman et al., 2016; Gobinath et al., 2018), Outliers were eliminated if two standard deviations above or below the mean and cytokine samples were eliminated if duplicates had high intra-assay coefficients of variations (>15%).

## 3. RESULTS

### 3.1 Maternal CORT treatment significantly lowered adrenal mass

CORT treatment significantly reduced relative adrenal mass compared to oil vehicle-treated animals (main effect of CORT; F(1, 33) = 103.120, *p* < 0.001, η_p_^2^ = 0.758; Figure 2A). No other significant effects or interactions were found (all *p*’s ≥ 0.403).

**Figure 2.**
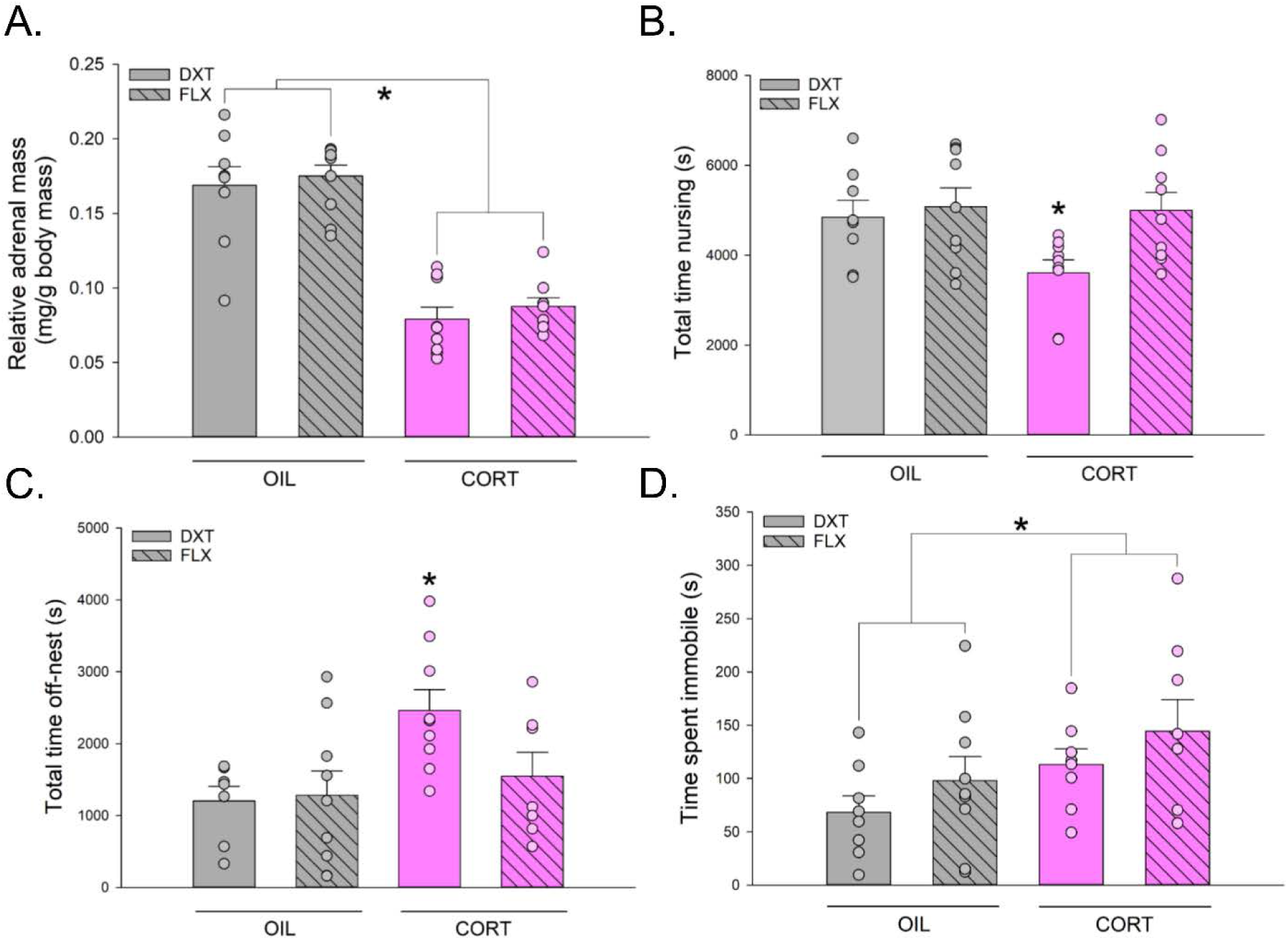
Effects of maternal corticosterone (CORT) and fluoxetine (FLX) treatment on relative adrenal mass, maternal care behaviour (total nursing, total off-nest behaviour), and immobility behaviour during forced swim test 2 (FST2), **A.** CORT treatment significantly reduced relative adrenal mass. *Indicate significance at *p* < 0.05. **B.** CORT + dextrose vehicle (DXT)-treated group showed significantly lower total nursing time compared to all other groups. *Indicate significance at *p* < 0.05. **C.** CORT+DXT treated groups showed a significantly higher total time off-nest when compared to all other groups. *Indicate significance at *p* < 0.05. **C.** CORT treatment significantly increased the time spent immobile in FST2. *Indicate significance at *p* < 0.05. Data represented in means + standard error of the mean (SEM), overlaid with individual data points.

### 3.2 Maternal CORT treatment reduced nursing and increased time off the nest and these effects were restored by FLX treatment

CORT+DXT showed significantly lower total time nursing compared to all other groups and postpartum FLX restored the CORT-induced decrease in nursing relative to OIL+DXT (a priori: *p* = 0.028, Cohen’s *d* = 1.148), OIL+FLX (*p* = 0.012, Cohen’s *d* = 1.330), and CORT+FLX (*p* = 0.012, Cohen’s *d* = 1.436); main effect of FLX: F(1, 31) = 4.713, *p* = 0.038, η_p_^2^ = 0.132, all other main or interaction effects were not significant (*p’s* > 0.09) Figure 2B).

CORT+DXT group showed significantly higher total time spent off-nest compared to OIL+DXT (a priori: *p* = 0.010, Cohen’s *d* = 1.756); OIL+FLX (*p* = 0.009, Cohen’s *d* = 1.249); CORT+FLX (*p* = 0.042, Cohen’s *d* =1.469); main effect of CORT, F(1, 28) = 6.207, *p* = 0.019, η_p_^2^ = 0.181; all other main or interaction effects were not significant (*p’s* > = 0.11 (Figure 2C)). No other significant effects on other maternal care behaviors were found (all *p*’s ≥ 0.126).

### 3.3 Maternal CORT treatment increased time spent immobile in FST2

Regardless of FLX, CORT treatment, significantly increased time spent immobile during FST2 compared to oil vehicle-treated animals (main effect of CORT, F(1, 29) = 4.464, *p* = 0.043, η_p_^2^ = 0.133 (Figure 2D)). No other significant effects of treatment on time spent immobile were found (all *p*’s ≥ 0.167).

There was a trend towards significance for CORT treatment to reduce time spent swimming compared to oil vehicle-treated animals (main effect of CORT, *p* = 0.086) but no other significant effects on time spent climbing were found (all *p*’s ≥ 0.379; data not shown). Time spent climbing was not affected by any of the treatments (all *p*’s ≥ 0.168).

### 3.4 Maternal FLX treatment increased IL-1β levels in the hippocampus; Maternal CORT treatment reduced two proinflammatory (TNF-α, IFN-γ) cytokine levels in the hippocampus

FLX treatment significantly increased levels of IL-1β in the hippocampus compared to vehicle-treated animals (main effect of FLX, F(1, 30) = 4.906, *p* = 0.034, η_p_^2^ = 0.140 (Figure 3A). FLX treatment did not significantly affect any other cytokines, chemokine, IL-6:IL-10 ratio, or IFN-γ:IL-10 ratio in the maternal hippocampus (all *p* ≥ 0.194).

**Figure 3.**
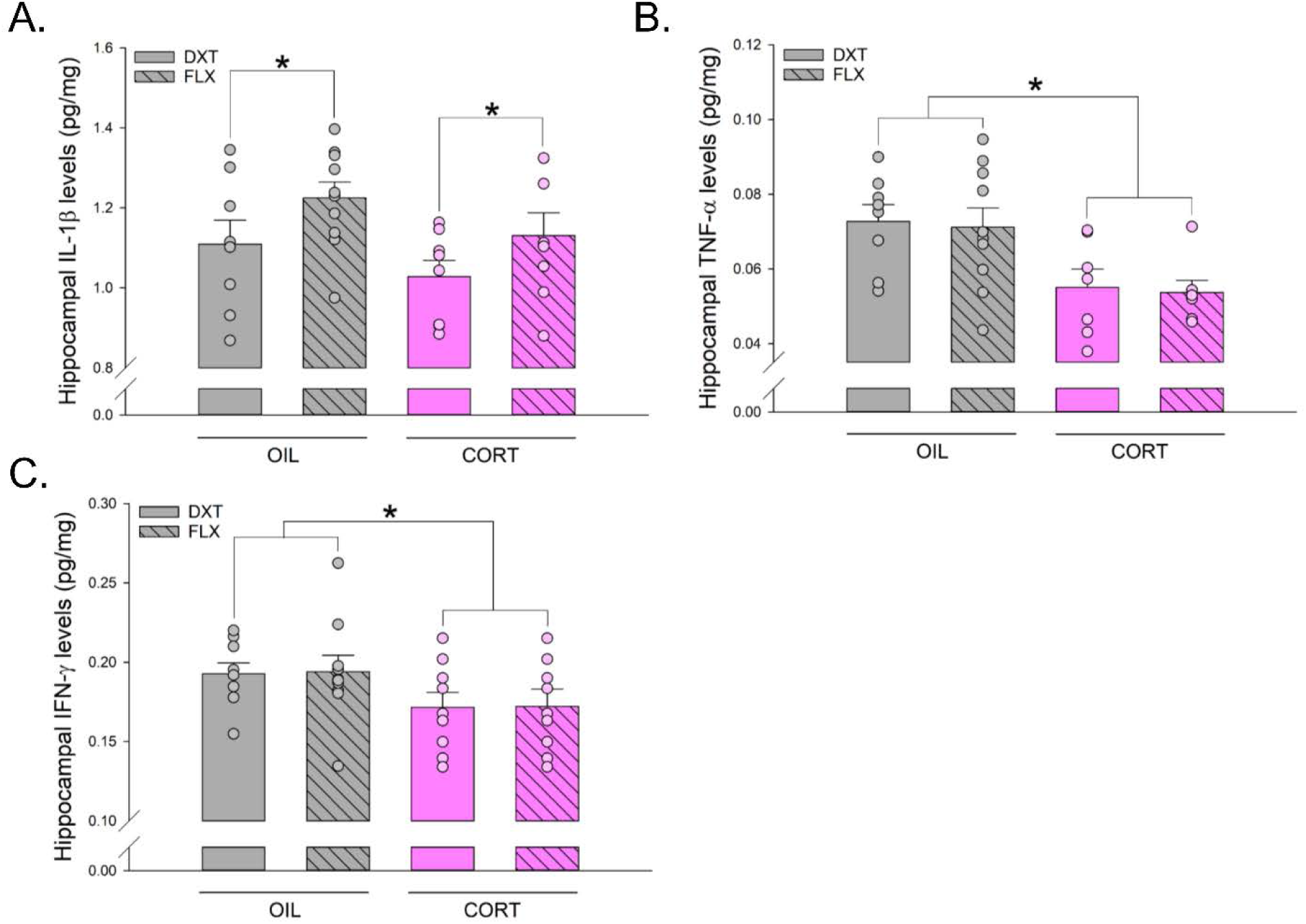
Effects of maternal corticosterone (CORT) and fluoxetine (FLX) treatment on hippocampal cytokine levels. **A.** FLX treatment significantly increased levels of hippocampal interleukin (IL)-1β. *Indicate significance at *p* < 0.05. **B.** CORT treatment significantly reduced hippocampal tumour necrosis factor(TNF)-α levels. *Indicate significance at *p* < 0.05. **C.** CORT treatment significantly reduced hippocampal interferon-gamma (IFN-γ) levels. *Indicate significance at *p* < 0.05. Data represented in means + standard error of the mean (SEM), overlaid with individual data points.

CORT treatment significantly reduced TNF-α and IFN-γ levels in the hippocampus compared to vehicle treatment (main effect of CORT: F(1, 28) = 13.771, *p* = 0.001, η_p_^2^ = 0.330; F(1, 31) = 5.108, *p* = 0.031, η_p_^2^ = 0.141, Figure 3 B and C, respectively). There were trends towards significance for an effect of CORT to reduce levels of IL-1β and IL-6 in the hippocampus compared to vehicle treatment, (*p* = 0.086, *p* = 0.067, respectively). CORT treatment did not significantly affect any other cytokines, chemokine, IL-6:IL-10 ratio, or IFN-γ:IL-10 hippocampus (all *p*’s ≥ 0.161; see Table 1). No other significant interaction effects between CORT and FLX treatment were observed in the maternal hippocampus (all *p* ≥ 0.339; Table 1). Further, there were no significant effects of CORT or FLX treatment, nor of their interaction, on cytokines, chemokines, IL6:IL-10 ratio, or IFN-γ:IL-10 levels in the maternal frontal cortex (all *p*’s ≥ 0.102; see Table 2).

**Table 1.** Two-way ANOVA results of hippocampal (HPC) cytokines and chemokine data. *Indicate significance at *p* < 0.05.

| **HPC** |  |  |  |  |
| --- | --- | --- | --- | --- |
| **Variable** |  | **F-value** | **p-value** | **$\eta_p^2$** |
| IFN-γ | CORT | 5.108 | *0.031 | 0.141 |
|  | FLX | 0.010 | 0.921 | < 0.001 |
| | CORT × FLX | 0.001 | 0.971 | < 0.001 |
| IL-10 | CORT | 2.073 | 0.161 | 0.069 |
|  | FLX | 0.319 | 0.576 | 0.011 |
| | CORT × FLX | 0.046 | 0.831 | 0.002 |
| IL-13 | CORT | 2.027 | 0.166 | 0.070 |
|  | FLX | 0.187 | 0.669 | 0.007 |
| | CORT × FLX | 0.947 | 0.339 | 0.034 |
| IL-1β | CORT | 3.159 | †0.086 | 0.095 |
|  | FLX | 4.906 | *0.034 | 0.140 |
| | CORT × FLX | 0.017 | 0.896 | < 0.001 |
| IL-4 | CORT | 1.187 | 0.285 | 0.038 |
|  | FLX | 0.529 | 0.473 | 0.017 |
| | CORT × FLX | 0.008 | 0.931 | < 0.001 |
| IL-5 | CORT | 0.050 | 0.825 | 0.002 |
|  | FLX | 1.778 | 0.194 | 0.064 |
| | CORT × FLX | 0.460 | 0.503 | 0.017 |
| IL-6 | CORT | 3.540 | †0.067 | 0.101 |
|  | FLX | 0.816 | 0.369 | 0.025 |
| | CORT × FLX | 0.260 | 0.610 | 0.008 |
| CXCL1 | CORT | 0.009 | 0.923 | < 0.001 |
|  | FLX | 0.454 | 0.505 | 0.014 |
| | CORT × FLX | 0.274 | 0.604 | 0.008 |
| TNF-α | CORT | 13.771 | *0.001 | 0.330 |
|  | FLX | 0.093 | 0.763 | 0.003 |
| | CORT × FLX | < 0.001 | 0.979 | < 0.001 |
| IL-6:IL-10 | CORT | 0.473 | 0.497 | 0.017 |
|  | FLX | 0.355 | 0.556 | 0.012 |
| | CORT × FLX | 0.063 | 0.803 | 0.002 |
| IFN-γ:IL-10 | CORT | 0.014 | 0.905 | < 0.001 |
|  | FLX | 0.066 | 0.800 | 0.002 |
| | CORT × FLX | 0.169 | 0.684 | 0.006 |

**Table 2.** Two-way ANOVA results of frontal cortex cytokines and chemokine data. *Indicate significance at *p* < 0.05.

| **Frontal cort** |  |  |  |  |
| --- | --- | --- | --- | --- |
| **Variable** | | **F-value** | **p-value** | $\eta_p^2$ |
| IFN-γ | CORT | 0.148 | 0.703 | 0.005 |
|  | FLX | 0.143 | 0.707 | 0.005 |
| | CORT × FLX | 0.420 | 0.522 | 0.013 |
| IL-10 | CORT | < 0.001 | 0.978 | < 0.001 |
|  | FLX | 0.305 | 0.585 | 0.010 |
| | CORT × FLX | 0.246 | 0.623 | 0.008 |
| IL-13 | CORT | 0.435 | 0.514 | 0.013 |
|  | FLX | 0.015 | 0.904 | < 0.001 |
| | CORT × FLX | 0.559 | 0.460 | 0.017 |
| IL-1β | CORT | 0.066 | 0.799 | 0.002 |
|  | FLX | 0.180 | 0.674 | 0.006 |
| | CORT × FLX | 0.124 | 0.727 | 0.004 |
| IL-4 | CORT | 0.006 | 0.938 | < 0.001 |
|  | FLX | 0.276 | 0.603 | 0.008 |
| | CORT × FLX | 0.378 | 0.543 | 0.011 |
| IL-5 | CORT | 0.034 | 0.855 | 0.001 |
|  | FLX | 0.288 | 0.585 | 0.009 |
| | CORT × FLX | 0.402 | 0.530 | 0.012 |
| IL-6 | CORT | 0.384 | 0.540 | 0.011 |
|  | FLX | < 0.001 | 0.987 | < 0.001 |
| | CORT × FLX | 1.139 | 0.294 | 0.033 |
| CXCL1 | CORT | 2.845 | 0.102 | 0.089 |
|  | FLX | 1.712 | 0.201 | 0.056 |
| | CORT × FLX | 0.004 | 0.952 | < 0.001 |
| TNF-α | CORT | 1.220 | 0.278 | 0.037 |
|  | FLX | 0.969 | 0.332 | 0.029 |
| | CORT × FLX | 0.012 | 0.912 | < 0.001 |
| IL-6:IL-10 | CORT | 0.057 | 0.812 | 0.002 |
|  | FLX | 0.123 | 0.728 | 0.004 |
| | CORT × FLX | 0.299 | 0.588 | 0.009 |
| IFN-γ:IL-10 | CORT | 0.043 | 0.837 | 0.001 |
|  | FLX | 0.004 | 0.949 | < 0.001 |
| | CORT × FLX | 0.345 | 0.562 | 0.012 |

### 3.5 Maternal fluoxetine treatment significantly reduced plasma 4’-pyridoxic acid, pyridoxal, and tryptophan concentrations, regardless of maternal corticosterone. Maternal corticosterone increased plasma pyridoxal and pyridoxal 5’-phosphate concentrations

Maternal FLX treatment significantly decreased plasma concentrations of tryptophan (Figure 4A), 4’-pyridoxic acid, and pyridoxal, (Figure 4B-C) compared to DXT-treated animals (main effect of FLX: F(1, 29) = 4.925, *p* = 0.034, η_p_^2^ = 0.145; F(1, 29) = 4.399, *p* = 0.044, η_p_^2^ = 0.132; F(1, 29) = 7.995, *p* = 0.008, η_p_^2^ = 0.216; respectively) but not pyridoxal 5’-phosphate (*p* = 0.546; Figure 4D). *A priori* analysis found that the OIL+FLX group showed significantly lower levels of plasma pyridoxal compared to all other groups (all *p*’s ≤ 0.006, all Cohen’s d ≥ 1.266; Figure 4C).

**Figure 4.**
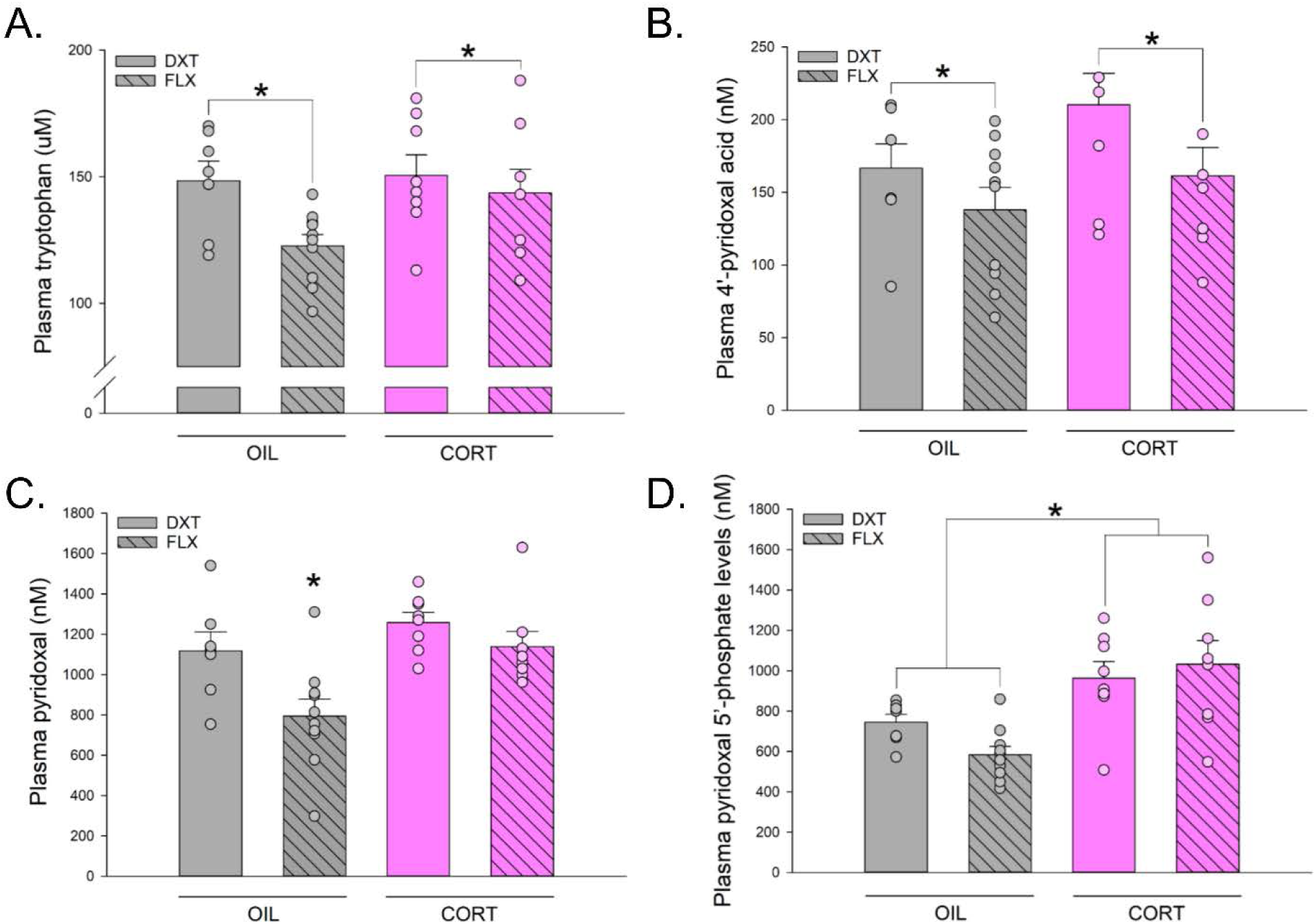
Effects of maternal corticosterone (CORT) and fluoxetine (FLX) treatment on plasma tryptophan levels, 4’-pyridoxic acid, pyridoxal levels, and pryidoxal 5’-phosphate. **A.** FLX treatment significantly reduced plasma tryptophan levels. *Indicate significance at *p* < 0.05. **B.** FLX treatment significantly reduced plasma 4’-pyridoxic acid levels. *Indicate significance at *p* < 0.05. **C.** OIL+FLX treatment significantly reduced pyridoxal levels comapred to all other groups. *Indicate significance at *p* < 0.05. **D.** CORT treatment significantly increased pyridoxal 5’-phosphate levels. *Indicate significance at *p* < 0.05. Data represented in means + standard error of the mean (SEM), overlaid with individual data points.

Maternal CORT significantly increased plasma pyridoxal (main effect of CORT, F(1, 29) = 9.549, *p* = 0.004, η_p_^2^ = 0.248; Figure 4C), and pyridoxal 5’-phosphate levels (main effect of CORT, F(1, 29) = 19.392, *p* ≤ 0.001, η_p_^2^ = 0.40; Figure 4D). Although CORT seemed to increase plasma levels of pyridoxic acid this was not statistically significant (*p* = 0.079; Figure 4C). There were no significant interaction effects for all metabolites (all *p’s* > 0.128).

### 3.6 Principal component analyses on brain cytokine and chemokine levels

Within the hippocampus, the PCA method found the first two principal components explaining 52.4% and 11.3% of the variation, respectively, together 63.70%. All nine cytokines and chemokines showed significant loadings onto the component 1 (all *p*’s < 0.001); IL-10 showed a significant negative loading onto component 2 (*p* < 0.001). CXCL1 (*p* = 0.004) and IL-1β (*p* < 0.001) showed a significant positive loading onto component 2 (Table 3).

**Table 3.** Component loadings for the hippocampal cytokine variance of the first two components identified using principal component analysis (PCA). Networks are interpreted using significant loadings set by variables denoted by *. All denoted values are determined by statistically significant contributions of variables towards the specific component (network). *Indicate significance at *p* = 0.001.

| Variables | Component 1 | Component 2 |
| --- | --- | --- |
| IFN-γ | *0.897 | 0.153 |
| IL-10 | *0.705 | *-0.550 |
| IL-13 | *0.809 | -0.268 |
| IL-1β | *0.660 | *0.572 |
| IL-4 | *0.780 | -0.122 |
| IL-5 | *0.642 | -0.159 |
| IL-6 | *0.576 | -0.141 |
| CXCL1 | *0.555 | *0.462 |
| TNF-α | *0.813 | 0.138 |

Within the frontal cortex, the PCA method found the first two principal components explaining 59.4% and 17% of the variation, respectively, together 72.39%. All cytokines and chemokines but IL-6 showed significant loadings on component 1 (all *p*’s < 0.001); IL-6 (*p* < 0.001) and CXCL1 (*p* < 0.001) showed a significant positive loading on component 2, IL-10 (*p* < 0.001) showed a significant negative loading on component 2 (Table 4).

**Table 4.**
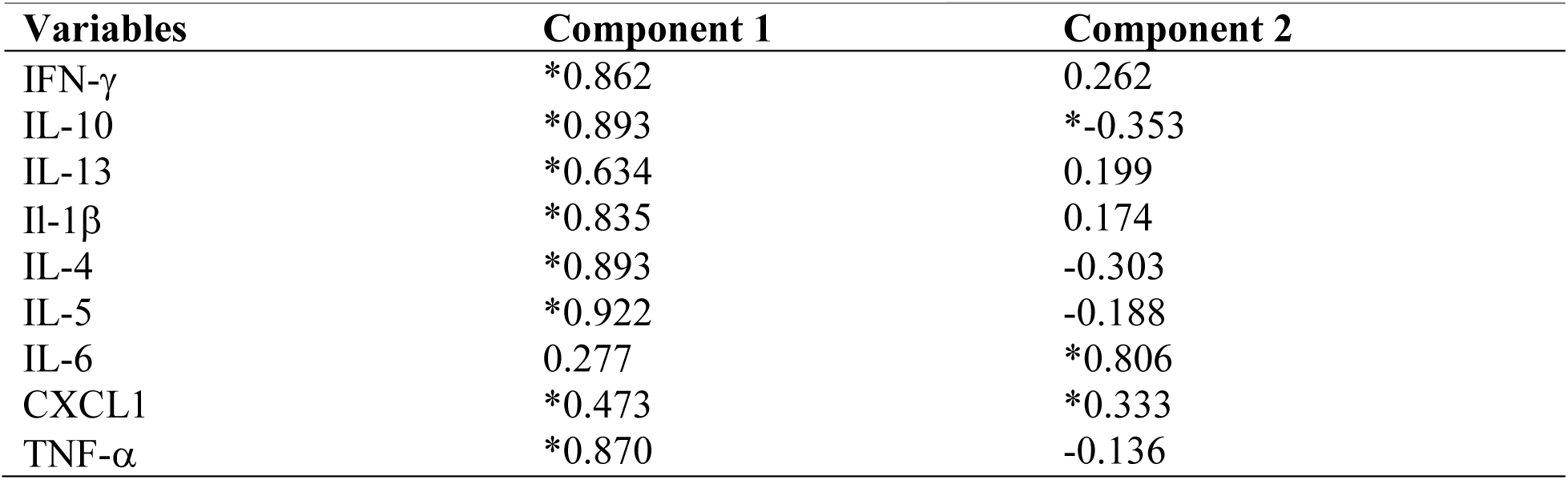
Component loadings for the frontal cortex cytokine variance of the first two components identified using principal component analysis (PCA). Networks are interpreted using significant loadings set by variables denoted by *. All denoted values are determined by statistically significant contributions of variables towards the specific component (network). *Indicate significance at *p* = 0.001

| Variables | Component 1 | Component 2 |
| --- | --- | --- |
| IFN-γ | *0.862 | 0.262 |
| IL-10 | *0.893 | *-0.353 |
| IL-13 | *0.634 | 0.199 |
| IL-1β | *0.835 | 0.174 |
| IL-4 | *0.893 | -0.303 |
| IL-5 | *0.922 | -0.188 |
| IL-6 | 0.277 | *0.806 |
| CXCL1 | *0.473 | *0.333 |
| TNF-α | *0.870 | -0.136 |

We analyzed sample scores onto principal component 1 using ANOVA, CORT treatment significantly decreased the sample scores onto principle component 1 within the HPC cytokine data set (main effect of CORT, F(1, 33) = 4.568, *p* = 0.040, η_p_^2^ = 0.122 (Figure 5A) but not the frontal cortex (p = 0.234; Figure 5B). No other significant main or interaction effects between CORT and FLX treatment were found when analyzing PCA sample scores within the HPC or frontal cortex cytokine data sets (all *p*’s ≥ 0.343). We did not find significant changes in samples scores onto principal component 2 using ANOVA in either HPC or frontal cortex data (all *p*’s ≥ 0.343).

**Figure 5.**
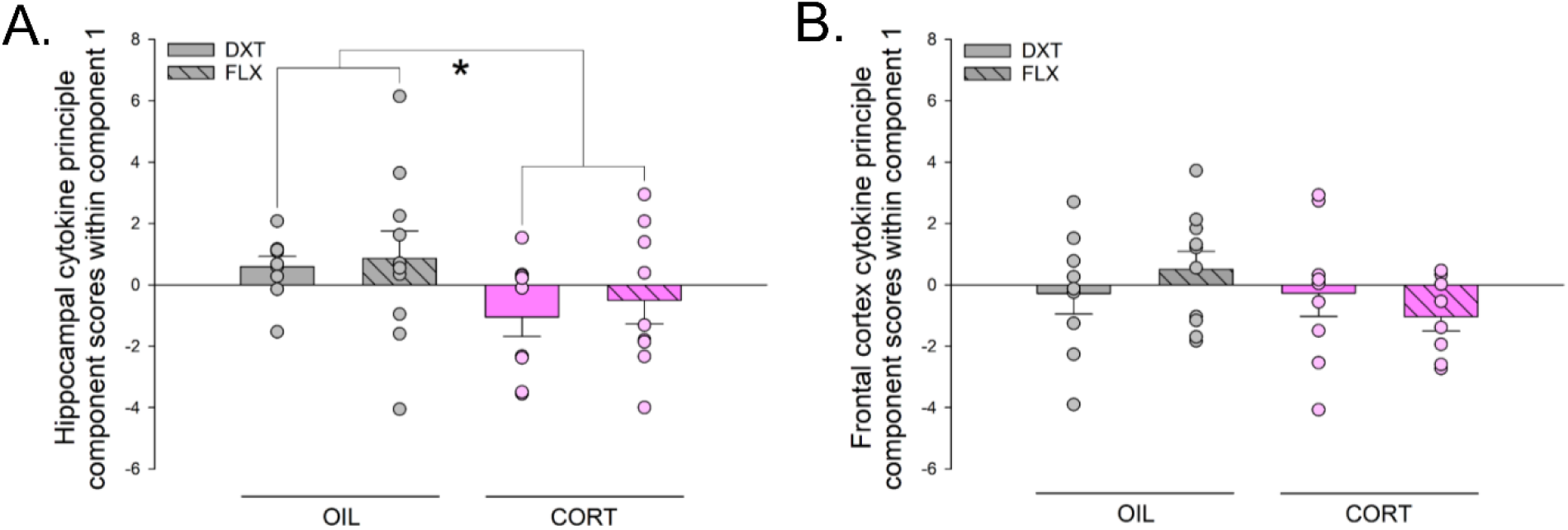
Effects of maternal corticosterone (CORT) and fluoxetine (FLX) treatment on hippocampal and frontal cortex cytokine PCA sample scores onto component 1. **A.** CORT treatment significantly lowered hippocampal cytokine sample scores onto component 1 in the principal component analysis. *Indicate significance at *p* < 0.05. **B.** CORT treatment significantly lowered frontal cortex cytokine sample scores onto component 1 in the principal component analysis. No statistically significant effects were found. Data represented in means + standard error of the mean (SEM), overlaid with individual data points.

## 4. DISCUSSION

Postpartum FLX treatment increased hippocampal IL-1β levels and reduced plasma tryptophan, 4’-pyridoxic acid, and pyridoxal concentrations regardless of CORT treatment. CORT treatment decreased two proinflammatory cytokines (TNF-α and IFN-γ) in the maternal hippocampus and increased plasma pyridoxal 5’-phosphate concentrations. Consistent with previous reports postpartum CORT treatment decreased maternal care behavior, increased passive-coping behavior, and reduced adrenal mass, whereas postpartum FLX restored maternal behaviors, but not rescue passive coping behaviors (Brummelte et al., 2006; Brummelte and Galea, 2010; Workman et al., 2016; Gobinath et al., 2018). Our findings point to the possibility that the lack of efficacy of FLX to restore passive coping is via its effects on IL-1β and tryptophan metabolism.

### 4.1 Corticosterone treatment-induced depressive-like behavior

Consistent with our previous research, postpartum CORT treatment significantly decreased total time spent nursing and increased total time spent off-nest (Brummelte et al., 2006; Brummelte and Galea, 2010; Workman et al., 2016; Gobinath et al., 2018). This finding is corroborated by other models of postpartum stress (Brunson et al., 2005; Ivy et al., 2008; Kurata et al., 2009; Nephew and Bridges, 2011; Moussaoui et al., 2016; Murgatroyd et al., 2015). As maternal care behavior is thought to be highly rewarding (Ferris et al., 2005; Mattson et al., 2001, 2003), the reduction in maternal care may be an expression of reduced reward-seeking or anhedonia-like behavior, which may be a representation of the clinical anhedonia phenotype of depression (Murgatroyd et al., 2015; reviewed by Post and Leuner, 2019).

We found that CORT treatment in the postpartum increased depressive-like behavior in the FST, consistent with previous reports (Brummelte et al., 2006; Brummelte and Galea, 2010; Workman et al., 2016; Gobinath et al., 2018), indicative of passive coping behavior. FST immobility has been interpreted as behavioral despair (reviewed in Krishnan and Nestler, 2011), which may be representative of the depressed mood and hopelessness clinical phenotype of depression. Overall, postpartum CORT injections induced behavioral endophenotypes of depression in rodent dams.

### 4.2 Postpartum corticosterone treatment reduced two proinflammatory cytokines in the hippocampus without affecting cytokine or chemokine levels in the frontal cortex

Patients with MDD (Howren et al., 2009; Dowlati et al., 2010; Haapakoski et al., 2015) and PND (Groer and Morgan, 2007; Osborne and Monk, 2013) show elevated levels of circulating proinflammatory cytokines, particularly TNF-α, IL-6, and IL-1β. Based on these clinical phenotypes, one might predict postpartum CORT treatment, which induces depressive-like behaviors (Brummelte et al., 2006; Brummelte and Galea, 2010; Workman et al., 2016; Gobinath et al., 2018) would elevate proinflammatory cytokines in our model. However, postpartum depression is associated with lowered peripheral IFN-γ levels compared to controls (Groer and Morgan, 2007), indicating the proinflammatory phenotype of PND is not seen. This effect is consistent with our findings that postpartum CORT treatment reduced two proinflammatory cytokines (TNF-α and IFN-γ) in the maternal hippocampus without significantly affecting other cytokines or chemokine measured in the hippocampus or the frontal cortex. Moreover, Groer and Morgan (2007) found significant changes in cytokine ratios (IFN-γ versus IL-10), suggesting possible changes in cytokine profiles. This effect is similar to what we found in our study, while we did not see changes in IFN-γ versus IL-10 ratio, we did see significant changes in cytokine profiles due to CORT treatment indicated by the PCA method. Interestingly, in a model of sub-chronic gestational stress (during gestational days 16-22), rat dams showed a decrease in IL-1β gene expression in the frontal cortex (Posillico and Schwarz, 2016). Although in the present study, we did not see any significant changes in the frontal cortex, we did see a non-significant reduction in hippocampal IL-1β protein with postpartum CORT.

Altogether, in our model of postpartum depression, we found reductions in two proinflammatory cytokines (TNF-α and IFN-γ). This finding is consistent with the anti-inflammatory effect of glucocorticoids in humans and rodents (reviewed by Sorrells and Sapolsky, 2007) and clinical data indicating lowered IFN-γ or no association between IL-6 and postpartum depression (Skalkidou et al., 2009; Okun et al., 2011) and PND (Corwin et al., 2008, 2015; Buglione-Corbett et al., 2018; Miller et al., 2019). These findings may suggest that perhaps with the altered immune profile of pregnancy and the postpartum, the changes in the inflammatory system in PND may not be representative of, or similar to, those found in MDD.

### 4.3 Fluoxetine treatment showed limited efficacy to reduce behavioral depressive-like endophenotypes

We found that FLX treatment rescued disrupted maternal care behavior early in the postpartum but not passive coping behavior later in the postpartum, in line with previous research (Brummelte et al., 2006, Brummelte and Galea, 2010, Workman et al., 2016, Gobinath et al., 2018). Interestingly, FLX treatment with exercise reduces passive-coping behavior in this model of postpartum depression (Gobinath et al., 2018). Other studies have also noted a lack of efficacy in the postpartum with pharmacological antidepressants (SSRIs) using coping behavior in the FST (Craft et al., 2010; Overgaard et al., 2018). Intriguingly, employing the hormone-withdrawal model of postpartum depression, neither the SSRI sertraline nor the tricyclic antidepressants, imipramine or desipramine, improved sucrose preference (Green et al., 2009; Navarre et al., 2010). Consistent with these findings, our data suggest a lack of efficacy of SSRIs to reduce depressive-like behavior in the postpartum period. It is interesting to note that when utilizing gestational stress to model depression onset during the antenatal period, the SSRI citalopram, was able to reduce passive-coping behavior (Haim et al., 2016). Thus, there appears to be a significant pathophysiological difference in SSRI efficacy and response to treatment depending on the etiology of PND in rodent models (gestational stress versus postpartum CORT exposure).

### 4.4 Fluoxetine treatment increased IL-1β in the hippocampus, increased plasma concentrations of tryptophan and altered vitamin B6 metabolism which may be related to the inefficacy of fluoxetine in the postpartum

FLX treatment increased levels of hippocampal IL-1β levels and decreased plasma tryptophan metabolism. Our results are consistent with another study that showed FLX-treated (non-partum) female mice had increased IL-1β gene expression in the hippocampus coupled with limited efficacy to reduce depressive-like behavior in both FST and the sucrose preference test (Duda et al., 2017). Overall, these data, coupled with our own, suggest an association between the lack of efficacy of FLX and increased IL-1β inflammation in the hippocampus of female rodents. Furthermore, SSRI treatment-resistant patients with MDD show elevated peripheral IL-1β levels after SSRI treatment (Syed et al., 2018), suggesting an association between elevated IL-1β levels and reduced treatment efficacy in both clinical studies and preclinical models. Future studies should determine if elevated IL-1β is causal and crucial for SSRI efficacy.

We also observed a reduction in plasma tryptophan concentrations in our rodent dams due to FLX treatment. This reduction may also contribute to the limited behavioral efficacy of maternal FLX in postpartum rats. Lower circulating concentration of tryptophan, a serotonin precursor, is observed in patients with MDD (Hughes et al., 2012; Myint et al., 2013; Ogawa et al., 2014; Doolin et al., 2018; Pu et al., 2020) and are thought to contribute towards a reduced rate of serotonin synthesis, leading to the onset of depressive symptom. Interestingly, in a study comparing tryptophan levels in patients with depression in the postpartum period to both health non-partum controls and healthy postpartum controls. Both healthy postpartum women and women with depression in the postpartum period had lower concentrations of peripheral tryptophan, but only the women endorsing depression in the postpartum period had lower concentrations of the tryptophan metabolite, kynurenine (Veen et al., 2016). This suggests that although parity itself can lower tryptophan concentrations and may lead to vulnerabilities to PND, those with PND show altered tryptophan-to-kynurenine metabolism compared to healthy non-partum and postpartum controls. Although in our study, we did not investigate any parity effects, future studies could explore if parity interacted with FLX treatment to influence tryptophan concentrations in the CORT-induced model of postpartum depression.

Additionally, we found significant decreases in vitamin B6 metabolism due to FLX treatment. Vitamin B6 is an essential nutrient that is not synthesized by the rodent and human body and appears in circulation as several different forms (reviewed in Parra et al., 2018). We found reductions in pyridoxal, an inactive form of vitamin B6 and a precursor of pyridoxal 5’-phosphate (the active form of vitamin B6), and 4’-pyridoxic acid, a secreted metabolite of pyridoxal 5-’phosphate. However, FLX treatment did not affect pyridoxal 5’-phosphate. Although low vitamin B6 derivatives are associated with depression (Hvas et al., 2004; Merete et al., 2008), this effect is controversial (Okereke et al., 2015). It should be noted that both plasma tryptophan and vitamin B6 concentrations can influence the brain (Troen et al., 2008; Schwarcz et al., 2012; Poulose et al., 2017). Vitamin B6 is an essential co-factor in tryptophan metabolism and a key methyl donor in homocysteine conversion. Elevated levels of homocysteine are also linked to depression (Bottiglieri et al., 2000; Sachdev et al., 2005). Thus, it is possible for vitamin B6 to play a significant modulatory role in depression onset. Here, the changes in vitamin B6 metabolites due to FLX treatment may be a representation of disrupted vitamin B6 metabolism. These effects may, in part, lower the efficacy of FLX to reduce depressive-like endophenotypes in postpartum rodent dams.

Interestingly, in human hippocampal progenitor cell cultures, the ability for the cytokine IL-1β to decrease neurogenesis is highly dependent on the tryptophan metabolic pathway (Zunszain et al., 2012). In addition to this association, vitamin B6 can inhibit cytokine (IL-1β, IL-6, and TNF-α) gene expression in cultured macrophages (Shan et al., 2020). Moreover, lower levels of peripheral pyridoxal 5’-phosphate concentrations correlate with high inflammation in both patients with rheumatoid arthritis and a rat model of rheumatoid arthritis (Chiang et al., 2005). Derivatives of vitamin B6 can also increase cell proliferation and the number of immature neurons in the hippocampus in adult male mice (Yoo et al., 2011). Overall, these data and ours together suggest the immune system, tryptophan metabolism, vitamin B6, and hippocampal integrity are highly correlated with each other and may act together to influence depression etiology and SSRI efficacy.

## 5. CONCLUSIONS

In conclusion, maternal CORT treatment-induced depressive-like behavior, while maternal FLX treatment showed limited efficacy to treat depressive-like behavior. Furthermore, our study identified two possible mechanisms that may contribute towards SSRI inefficacy in the postpartum: (1) maternal FLX increased hippocampal IL-1β levels and (2) reduced plasma levels of tryptophan and vitamin B6 metabolism. Ultimately, our study contributes to the body of literature to suggest a possible association between an altered cytokine profile and the pathophysiology of PND. Future studies are needed to determine if altered immune profiles, tryptophan and vitamin B6 metabolism can act as biomarkers and improve pharmacological treatments for PND.

## ACKNOWLEDGEMENT

The authors would like to thank all the wonderful staff members at the Centre for Disease Modelling animal facility at the University of British Columbia for the continuous support and help throughout the project. Financial support for this research was provided by an operating grant from the Canadian Institutes for Health Research (MOP142308) to LAMG. WQ is supported by a 4-year Fellowship granted by the University of British Columbia, Faculty of Medicine, Graduate Program in Neuroscience, and the University of British Columbia Institute of Mental Health Marshalls Scholars Program in Mental Health, Department of Psychiatry.

